# The effects of predator odor (TMT) exposure and mGlu_3_ NAM pretreatment on lasting behavioral and molecular adaptations in the insular cortex and BNST

**DOI:** 10.1101/2021.05.07.443122

**Authors:** Ryan E. Tyler, Maya N. Bluitt, Julie L. Engers, Craig W. Lindsley, Joyce Besheer

## Abstract

A stressor can trigger adaptations that contribute to neuropsychiatric disorders. Predator odor (TMT) exposure is an innate stressor that produces lasting adaptations. TMT exposure may activate metabotropic glutamate receptor 3 (mGlu_3_), triggering excitatory corticolimbic adaptations that underlie behavioral changes. To evaluate functional involvement, the mGlu_3_ negative allosteric modulator (NAM, VU6010572; 3 mg/kg, i.p.) was administered before TMT exposure in male, Long Evans rats. Two weeks after stressor, rats underwent behavioral testing (context re-exposure, zero maze and acoustic startle response) followed by RT-PCR gene expression in the insular cortex and BNST. During the TMT exposure, rats displayed stress-reactive behaviors that were not affected by the VU6010572. During the context re-exposure, prior TMT exposure and VU6010572 pretreatment both produced a hyperactive response. TMT exposure did not affect zero maze or ASR measures, but VU6010572 increased time spent in the open arms and habituation to ASR, indicating anxiolytic-like effects. In the insular cortex, TMT exposure resulted in excitatory adaptations as shown by increased expression of mGlu (*Grm3, Grm5*), NMDA (*GriN2A, GriN2B, GriN2C, GriN3A, GriN3B*) and AMPA (*GriA3*) receptor transcripts. Interestingly, mGlu_3_ signaling during stressor mediated *GriN3B* upregulation. Stress reactivity during TMT exposure was associated with *Grm5, GriN2A, GriN2C*, and *GriA3* upregulation in the insular cortex and context re-exposure reactivity in the TMT/vehicle, but not the TMT/mGlu_3_ NAM group. In the BNST, *GriN2A, GriN2B* and *GriN3B* were increased by VU6010572, but TMT prevented these effects. These data demonstrate that mGlu_3_ signaling contributes to the lasting behavioral and molecular adaptations of predator odor stressor.

## 1. Introduction

Stress-related neuropsychiatric disorders are associated with dysregulated glutamatergic function [1-6]. Stress directly alters glutamatergic neurotransmission through glucocorticoid-receptor (GR) binding on glutamate neurons [7]. For example, stress models (e.g., forced-swim, tail-suspension, foot shock, glucocorticoid treatment) have repeatedly been shown to acutely increase presynaptic glutamate release in the brain [7-14]. Further, acute stressor exposure (e.g., forced-swim, tail suspension, elevated-plus maze) potentiates *N*-methyl-D-aspartate receptor (NMDA) and α-amino-3-hydroxy-5-methyl-4-isoxazolepropionic acid (AMPA) receptor currents and surface expression through a GR-dependent mechanism on medial prefrontal cortex (mPFC) pyramidal neurons [15]. Overall, stress has been shown to have a multitude of acute and lasting effects specifically on cortico-limbic glutamate function that potentially contribute to maladaptive responses that increase vulnerability for the development of neuropsychiatric disorders[7, 16, 17].

Metabotropic glutamate (mGlu) receptors are potential therapeutic targets for many neuropsychiatric disorders [1-3, 6, 18], and modulate excitatory cell function through intracellular cascade events that can result in lasting adaptations to the cell in response to stressors [6, 19]. Group II mGlu receptors (mGlu_2_ and mGlu_3_) are coupled to Gi-proteins that reduce cellular activity as a result of activation [6]. Traditionally, Group II mGlu receptors are considered predominantly located on presynaptic terminals and their activation attenuates presynaptic neurotransmitter release [6]. However, mGlu_3_ receptors have been found on both pre- and post-synaptic neurons, as well as astrocyte projections in the brain [20, 21]. Regardless of localization, mGlu_2/3_ receptor blockade has consistently been shown to have antidepressant- and antianxiety-like behavioral effects in rodents induced by stress with overlapping brain mechanisms to that of ketamine and other glutamate-targeting therapeutics [16, 17, 22-27]. Therefore, mGlu_2/3_ receptor blockade has been proposed as an alternative to ketamine for its therapeutic potential without the psychogenetic and addiction-liability risks associated with ketamine [17]. The mechanisms by which mGlu_2_ and mGlu_3_ signaling exert their therapeutic effects are important for understanding novel mechanisms of glutamate-targeting antidepressant-like effects. However, prior to the recent development of highly specific negative allosteric modulators (NAMs) for mGlu_3_ receptors [28, 29], the specific role of mGlu_3_ vs. mGlu_2_ receptors in these effects had been elusive. The same drug class of mGlu_3_ NAMs that were used in these experiments (VU6010572, first generation: VU0650786) have recently been employed to show that mGlu_3_ mediates a form of synaptic plasticity in cortico-limbic circuitry that is sensitive to stress [19, 20, 27, 28] and exhibits rapid anti-depressant-like effects [27]. Restraint stress before electrophysiological measurements occluded mGlu_3_-mediated long-term depression (LTD) in a manner consistent with mGlu_3_ loss of function. Prior studies show that blocking mGlu_3_ activation *in vivo* prior to restraint stress restored mGlu_3_-LTD [19]. As previously mentioned, mGlu_2/3_ receptor blockade reverses stress-induced depressive-like phenotypes in rodents [17]. Therefore, these previous findings suggest that synaptic plasticity induced by mGlu_3_-signaling during a stress event may play a role in the lasting behavioral and glutamatergic adaptations of stressor exposure.

Cortico-limbic circuitry is implicated in stress-related neuropsychiatric disorders. In animal models, the insular cortex and the bed nucleus of the stria terminalis (BNST) both play an important role in stress responses [30, 31]. The insular cortex is commonly associated with the integration of sensory and interoceptive processing [32, 33], which is highly sensitive to stress and anxiety in preclinical models [34, 35] and implicated in the etiology of neuropsychiatric disorders [36]. For example, inactivation of the anterior insular cortex produced anxiogenic-like effects, whereas activation of the insular cortex produced anxiolytic-like effects as assessed by the elevated plus maze test [34]. Clinically, anticipation of unpredictable stress increased BOLD signal in the insular cortex as assessed by fMRI in human subjects [31]. The BNST is also associated with stress-related neuropsychiatric disorders [37] and animals models of stress adaptations [38]. For example, restraint stress in mice increased c-Fos immunoreactivity in CRF-expressing neurons of the dorsal BNST [39]. Additionally, inactivation of the BNST during predator odor exposure stress blocked stressor-induced freezing behavior in rodents [40], indicating involvement in the fear-like response to TMT. Finally, the insular cortex and BNST show reciprocal projections that are likely involved in the lasting plasticity from a stress response [41]. Therefore, the focus of the present work is on glutamatergic gene expression adaptations following stressor exposure in the insular cortex and BNST. Genes that encode for glutamate receptors were evaluated: AMPA: *GriA1, GriA2, GriA3*, NMDA: *GriN1, GriN2A, GriN2B, GriN2C, GriN2D, GriN3A GriN3B*, and mGlu: *Grm2, Grm3*, and *Grm5*.

Animal models have used exposure to the scent of a predator as a stressor that produces lasting behavioral consequences with relevance to stress-related neuropsychiatric disorders [42-55]. For example, exposure to the synthetically derived fox odor 2,5□dihydro□2,4,5□trimethylthiazoline (TMT) produces avoidance and freezing behaviors, indicative of a fear□like response and increased serum corticosterone, reflecting a stress response [47, 50, 56, 57]. Other studies have showed lasting anxiety□like and hyperarousal behavior, as well as avoidance of a TMT□paired context or cue [43, 51]. Our lab recently showed that TMT exposure also produces lasting adaptations (4 weeks after TMT) in glutamate-related gene expression (*Grm2, Grm5, Grm7, Shank3, Homer1, Slc1A3*) in corticolimbic brain regions [58]. Of relevance to the present work, we have reported decreased *Grm3* gene expression 2 days after TMT exposure [58]. Given recent physiology findings previously discussed [29], we hypothesized that this change may reflect a transcriptional mechanism for loss of function following stressor-induced activation of mGlu_3_. Furthermore, mGlu_3_ receptor activation during stressor may be involved in lasting adaptations to glutamatergic cortico-limbic circuitry [16, 17, 23-26]. Therefore, the present study blocks mGlu_3_ signaling during the TMT exposure to evaluate the functional involvement of mGlu_3_ activation during a stressor.

Therefore, the goal of the present work was to block mGlu_3_ receptor activation (using VU6010572) *during* the TMT stressor and evaluate its functional implications on the *lasting* adaptations as a consequence of stressor. To capture the *lasting* behavioral and molecular consequences of the TMT stressor, rats remained undisturbed in the home cage for 2 weeks (operationally defined here as an “incubation period”) before behavioral testing (context re-exposure, zero maze, and acoustic startle response) and molecular assessments (gene expression for glutamate receptor transcripts) were conducted. Therefore, these experiments provide insight into the role of mGlu_3_-signaling *during* a stressor on *lasting* stressor-induced adaptations, and in doing so also evaluate the potential long-term therapeutic effects of mGlu_3_ NAM (VU6010572). These data could provide us with greater mechanistic insight into how stressor may influence vulnerability to the development of a neuropsychiatric disorder.

## 2. Methods

### 2.1 Animals

Male Long□Evans rats (n□=□56; Envigo, Indianapolis, IN) were used for all experiments. Rats arrived at 7□weeks and were handled for at least 1□minute daily for 1□week prior to experiments. To obviate potential stress-transfer effects, all rats arrived and remained single-housed throughout the duration of experiments, consistent with our previous publications [58]. Rats were housed in ventilated cages with access to food and water ad libitum and maintained in a temperature and humidity-controlled vivarium with a 12-hour light/dark cycle. All experiments were conducted during the light cycle. Rats were under continuous care and monitoring by veterinary staff from the Division of Comparative Medicine at UNC□Chapel Hill. All procedures were conducted in accordance with the NIH Guide to Care and Use of Laboratory Animals and institutional guidelines.

### 2.2 Compounds

VU6010572 was provided by Dr. Craig Lindsley and synthesized according to [29]. VU6010572 was dissolved in 45% β-cyclodextrin. A 1.5 mg/mL solution was made and injected at a volume of 2 mL/kg intraperitoneally (i.p.) to achieve a 3 mg/kg dose. An equal volume of 45% β-cyclodextrin was used as vehicle. The predator odor used was 2,5□dihydro□2,4,5□trimethylthiazoline (TMT; 97% purity; SRQ Bio, Sarasota, FL).

### 2.3 Assessment of mGlu_3_ NAM in open field test

Rats were transported in the home cage to the behavioral testing room and allowed to acclimate for 30 min before starting the experiment. VU6010572 (0, 1, 3, and 6 mg/kg; n=4/dose) was administered 45 min prior to the open field test to examine locomotor effects. Rats were placed in the center of the open field (43 cm × 43 cm; Med Associates, St. Albans, VT) enclosed in a sound attenuating cubicle with the lights off for a 60-min test. Locomotor activity was assessed by 32 orthogonal infrared beams and quantified using Activity Monitor software (Med Associates).

### 2.4 Assessment of mGlu_3_ NAM pretreatment on TMT exposure

Rats (n=12/grp) were injected with VU6010572 (0 or 3 mg/kg, i.p.) and returned to the home cage for 45 mins prior to TMT-exposure testing. TMT exposure test chambers and experimental set-up were identical to those used in [56, 58]. Rats were transported from the vivarium in the home cage to a separate, well□ventilated room that contained the test chambers in which rats were exposed to TMT (45.72□×□17.78□×□21.59□cm; UNC Instrument Shop, Chapel Hill, NC). Only one rat was placed in each chamber. The length of the back wall of the test chambers was opaque white with two opaque black side walls and a clear, plexiglass front wall to enable video recordings and a clear sliding lid. A small, metal basket was hung on the right□side wall (17.8 cm above the floor) to hold a piece of filter paper. 10 μL of TMT (2,5□dihydro□2,4,5□trimethylthiazoline) or water for controls was pipetted onto the filter paper in the metal basket immediately prior to putting the rat in the chamber. The control group was always run before the TMT group to prevent odor contamination. The odor exposure session lasted 15 mins and was video recorded for evaluation of behavior using ANY□maze Video Tracking System (Version 6.12, Stoelting Co. Wood Dale, IL). After TMT exposure, rats were returned to the home cage and remained undisturbed in the home cage for 2 weeks as an incubation period prior to starting behavioral assessments. Two rats that were initially assigned to the TMT/mGlu_3_ NAM group were moved to the TMT/vehicle group prior to injection due to unintentional loss of mGlu_3_ NAM solution. One other animal in the TMT/mGlu_3_ NAM group was removed from the study due to a failed i.p. injection. Final sample sizes: (n=12 - ctrl/vehicle; n=12 - ctrl/mGlu_3_ NAM; n=14 - TMT/vehicle; n=9 - TMT/mGlu_3_ NAM).

### 2.5 Context Re-exposure

Two weeks following TMT exposure, rats were returned to the same test chambers in which they had been previously exposed to TMT. No TMT was used for this context re-exposure test. Identical to the TMT exposure, this test lasted 15 min in duration and was video recorded for behavioral assessments identical to the TMT exposure behavioral measures.

### 2.6 Elevated Zero Maze

24 hours after the context re-exposure test (15 days post-TMT exposure), rats were evaluated on an elevated zero maze. Rats were transported in their home cages to the behavioral testing room and allowed to acclimate for 30 min prior to testing. The zero maze was made up of a circular platform with a diameter of 99 cm raised above the floor to a height of approximately 70 cm. The maze was divided equally into four quadrants/arms. Two enclosed arms contained two walls with one front wall [33.02 cm (H)] and back wall [64.52 cm (H)]. The exposed arms were located on opposite sides of the circular platform and were bordered with approximately 5 cm rim to prevent the rat from falling off the circular platform. Rats were placed in one open arm at the start of the 5 min test. These experiments were video recorded for analysis.

### 2.7 Acoustic Startle Response

24-hours after the zero-maze test (16 days pots-TMT exposure), rats were evaluated on an acoustic startle response test using an acoustic startle response system (S-R Lab; San Diego Instruments, San Diego, CA). Rats were transported in their home cages to the behavioral testing room and allowed to acclimate for 30 min prior to testing. Rats were placed in a cylinder Plexiglas animal enclosure located within a sound-attenuating test chamber that included an exhaust fan, a sound source, and an internal light that was turned off during the test. At the start of each test, rats underwent a 5-min habituation period during which 60 dB of background white noise was present. The background noise was present during the entire test session. The test session consisted of 30 trials of a 100 ms burst of a 110 dB startle. Each trial was separated by a 30 to 45-s randomized intertrial interval. Startle response (amplitude) was measured with a high-accuracy accelerometer mounted under the animal enclosure and analyzed with SR-Lab software.

### 2.8 Brain tissue collection and sectioning

24 hours after the acoustic startle test (17 days post-TMT exposure), a sub-set of rats (n=5 control/vehicle, n=5 control/mGlu_3_ NAM, n=12 TMT/vehicle, n=9 TMT/mGlu_3_ NAM) were sacrificed for qPCR analysis. More rats were allocated to the TMT group because previous studies showed that TMT exposure can produce distinct sub-groups within the TMT-exposed group that differ in brain gene expression [51]. Rats were anesthetized in 5% isoflurane immediately prior to brain extraction. Brains were rapidly extracted, and flash frozen with isopentane (Sigma□Aldrich, MI), then stored at −80 °C until brain region sectioning. Brains were sectioned on a cryostat (−20 °C) up to a predetermined bregma for each region of interest (ROI) according to [59]. Then, a micropunch tool was used to remove tissue specific to each brain region as illustrated in Table 1. Brain sections were stored at −80°C until qPCR analysis. *qPCR Analysis*

**Table 1 –.**
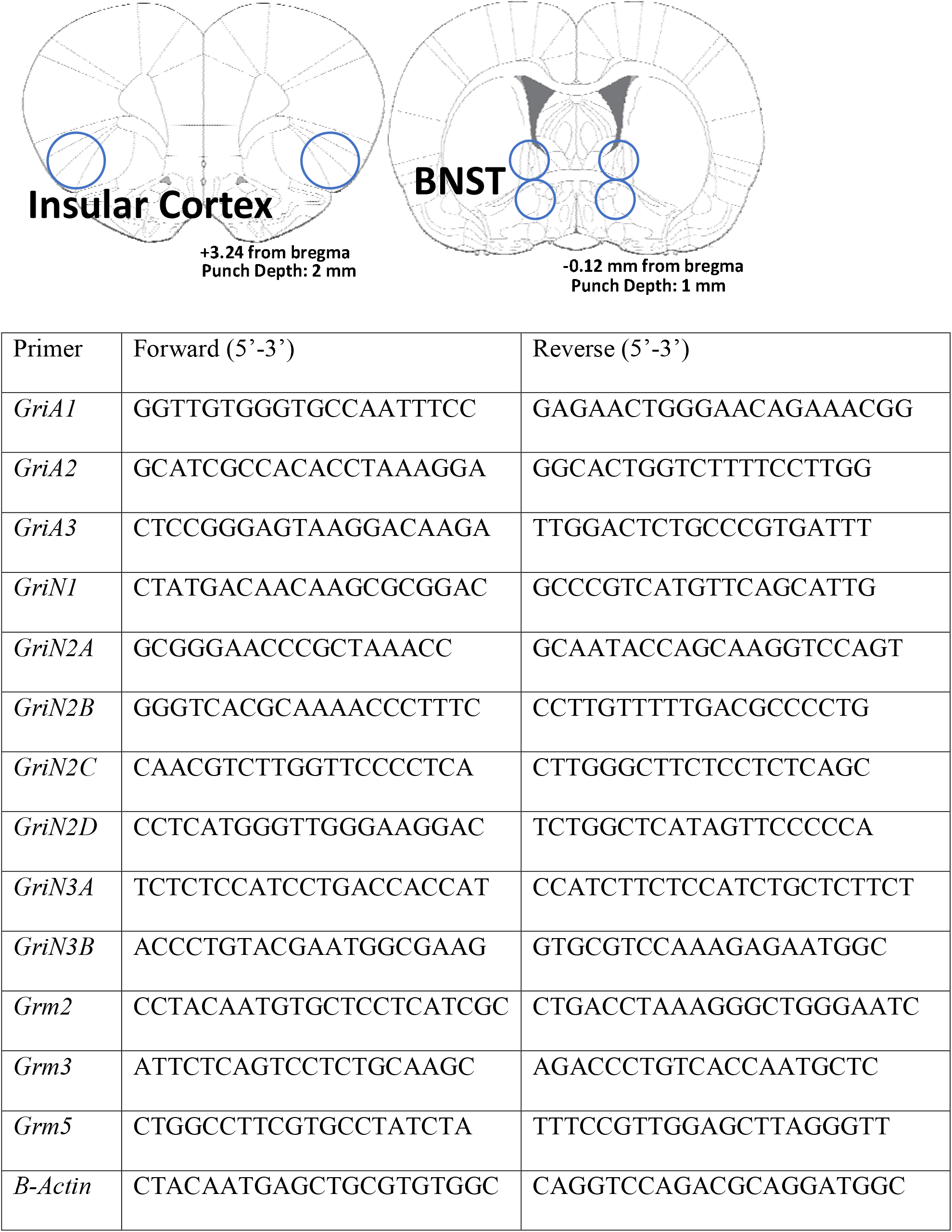
qPCR Tissue extraction and Primer Sequences.

### 2.9 RNA Extraction□

RNA was extracted from brain tissue using the RNeasy Mini Kit (Qiagen, Venlo, Netherlands) according to the manufacturer’s instructions. RLT lysis buffer containing β□mercaptoethanol (Sigma Aldrich) was used for tissue homogenization. RNA concentration and purity for each sample were determined using a Spectrophotometer (Nanodrop 2000, ThermoScientific).

### 2.10 Reverse Transcription□

RNA was reverse transcribed into cDNA using the QuantiNova Reverse Transcription Kit (Qiagen, Venlo, Netherlands) according to the manufacturer’s instructions. Following reverse transcription, all samples were diluted 1:10 with water and stored at −20°C before RT□PCR experiments.

### 2.11 RT□PCR□

The QuantStudio3 PCR machine (ThermoFisher) was used for all experiments. Using a 96□well plate, each sample was run in triplicate using 10 μL total volume per well with the following components: PowerUp Syber green dye (ThermoFisher, containing ROX dye for passive reference), forward and reverse primers (Eton Biosciences Inc., NC) and cDNA template. The PCR was run with an initial activation for 10 minutes at 95 °C, followed by 40□cycles of the following: denaturation (95°C for 15□seconds), annealing/extension (60°C for 60□seconds). Melt curves were obtained for all experiments to verify synthesis of a single amplicon. All primers targeted genes that encode for glutamate receptors and include: *GriA1, GriA2, GriA3, GriN1, GriN2A, GriN2B, GriN2C, GriN2D, GriN3A GriN3B, Grm2, Grm3*, and *Grm5*. All primer sequences are displayed in Table 1.

## 3. Data Analysis

### 3.1 Open Field Test

Locomotion was quantified over time in 10-min bins and analyzed using a two-way ANOVA with time as a repeating factor. A one-way ANOVA was used to compare total locomotion between doses.

### 3.2 TMT Exposure and Context Re-exposure

Analysis of TMT exposure and context re-exposure behaviors are identical to {Tyler, 2020 #49}. Using ANY□maze, the length of the rectangular TMT exposure chamber was divided into two compartments for analysis (TMT side and non□TMT side). The basket containing TMT was located on the far end of the TMT side. Dependent measures assessed via ANY-maze included time spent immobile, time spent of the TMT side of the chamber, distance traveled and midline crossings (the number of times the animal crossed between the TMT and non-TMT side). Immobility was operationally defined as the absence of movement other than respiration for longer than 2 seconds, and also captures freezing behavior during the TMT exposure [58]. Time spent digging in the bedding (see [56, 58]) and time spent grooming were quantified manually by an experimenter blind to experimental conditions. For the context re-exposure, two rats were removed from analysis due to a camera recording error. Context re-exposure sample sizes: (n=11 - control/vehicle; n=12 - control/mGlu_3_ NAM; n=13 - TMT/vehicle; n=9 - TMT/mGlu_3_ NAM).

These dependent measures were analyzed by two□way ANOVAs and Sidak’s multiple comparison test was used for all post hoc analyses. Pearson correlational analyses were conducted between stress-reactive behaviors to the TMT exposure and to the context re-exposure reactivity in the TMT/Vehicle group. Positive correlations identified in the TMT/Vehicle group were evaluated in all other experimental groups.

### 3.3 Zero Maze

The time spent in the open arms was quantified using ANY-maze software and analyzed by two□way ANOVA with Sidak’s multiple comparison test for post hoc analyses. Three rats were excluded from this analysis – two due to falling off of the zero maze, and another was determined to be a statistically significant outlier. Sample sizes: (n=10 - control/vehicle; n=12 - control/mGlu_3_ NAM; n=13 - TMT/vehicle; n=9 - TMT/mGlu_3_ NAM).

### 3.4 Acoustic Startle Response

The peak startle response (amplitude) was averaged for all 30-trials. In order to assess how startle response changes over time, and to evaluate habituation, we averaged the first and last 15-trials. Next, we calculated the percent change between the average of the first 15 trials and last 15 trials as follows:

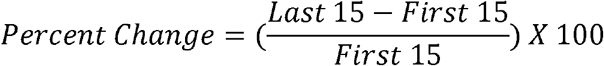

This measure was defined as “Habituation.” Two rats were removed from this analysis due to an experimental error. Sample sizes: (n=11 - control/vehicle; n=11 - control/mGlu_3_ NAM; n=13 - TMT/vehicle; n=9 - TMT/mGlu_3_ NAM). Data were analyzed by two□way ANOVA with Sidak’s multiple comparison test for post hoc analyses.

### 3.5 Gene expression assessments

The ΔΔCt method was used to determine fold change relative to controls [60]. Fold changes were normalized so that average control fold change equaled 1. Samples sizes for gene expression analysis started as: n=5 control/vehicle; n=5 control/mGlu_3_ NAM; n=12 TMT/vehicle; n=9 TMT/mGlu_3_ NAM (see methods). Three rats were removed from analysis due to brain extraction errors. Sample size for the insular cortex: n=4 control/vehicle; n=5 control/mGlu_3_ NAM n=12 TMT/vehicle; n=7 TMT/ mGlu_3_ NAM. For *Grm3* in the insular cortex, one additional rat was removed from analysis as a statistically significant outlier in the control/vehicle group. For the BNST, 3 additional samples were lost as a result of an RNA isolation errors, resulting in n=4 - control/vehicle; n=4 control/mGlu_3_ NAM; n=10 TMT/vehicle; n=7 TMT/mGlu_3_ NAM. Data were analyzed using 2-way ANOVAs, with Sidak’s multiple comparison test for post hoc analyses. All gene expression data are reported as mean fold□change ±□SEM. For all analyses, significance was set at P□≤□0.05.

## 4. Results

### 4.1 mGlu_3_ NAM (VU6010572) dose testing with open field

Because this VU6010572 compound has not been tested in rats, we used an open-field test to determine the appropriate dose for our studies. The two-way RM ANOVA showed no effect of the mGlu_3_ NAM dose range (0, 1, 3, 6, mg/kg, IP) on distance traveled over time, but did show a main effect of time (F (1.6, 14.5) = 42.81, p < 0.0001), with decreased distance traveled over time. Total distance traveled also showed no effect of VU6010572 dose (cm; 0 mg/kg: 4045 ± 843.6; 1 mg/kg: 3758 ± 802.2; 3 mg/kg: 2675 ± 632.3; 6 mg/kg: 1776 ± 523.1), however, visual inspection of 6 mg/kg VU6010572 showed a trend for decreased locomotion in the open field. Therefore, 3 mg/kg was chosen as the drug dose for TMT exposure experiments, consistent with dosing in mice [29].

### 4.2 TMT exposure produces stress-reactive behaviors that are not affected by mGlu_3_ NAM

Figure 1 A-F shows the behavioral responses during TMT exposure. All behavioral measures (e.g., digging, immobility, time spent on the TMT side, grooming, distance traveled and midline crossings) showed a main effect of TMT exposure and no significant effect of mGlu_3_ NAM or interaction. TMT exposure increased time spent digging (Fig. 1A, F (1, 43) = 38.32, p < 0.0001) and time spent immobile (Fig. 1B, F (1, 43) = 70.96, p < 0.0001). TMT exposure decreased time spent on the TMT side of the test chamber (Fig. 1C, F (1, 43) = 61.55, p < 0.0001) and time spent grooming (Fig. 1D, F (1, 43) = 6.02, p = 0.01). Distance traveled (Fig. 1E, F (1, 43) = 7.15, p = 0.01) and midline crossings (Fig. 1F, F (1, 43) = 10.44, p = 0.002) were decreased as a main effect of TMT exposure. These results demonstrate behavioral stress-reactivity to TMT exposure that is not affected by pretreatment with the mGlu_3_ NAM.

**Figure 1.**
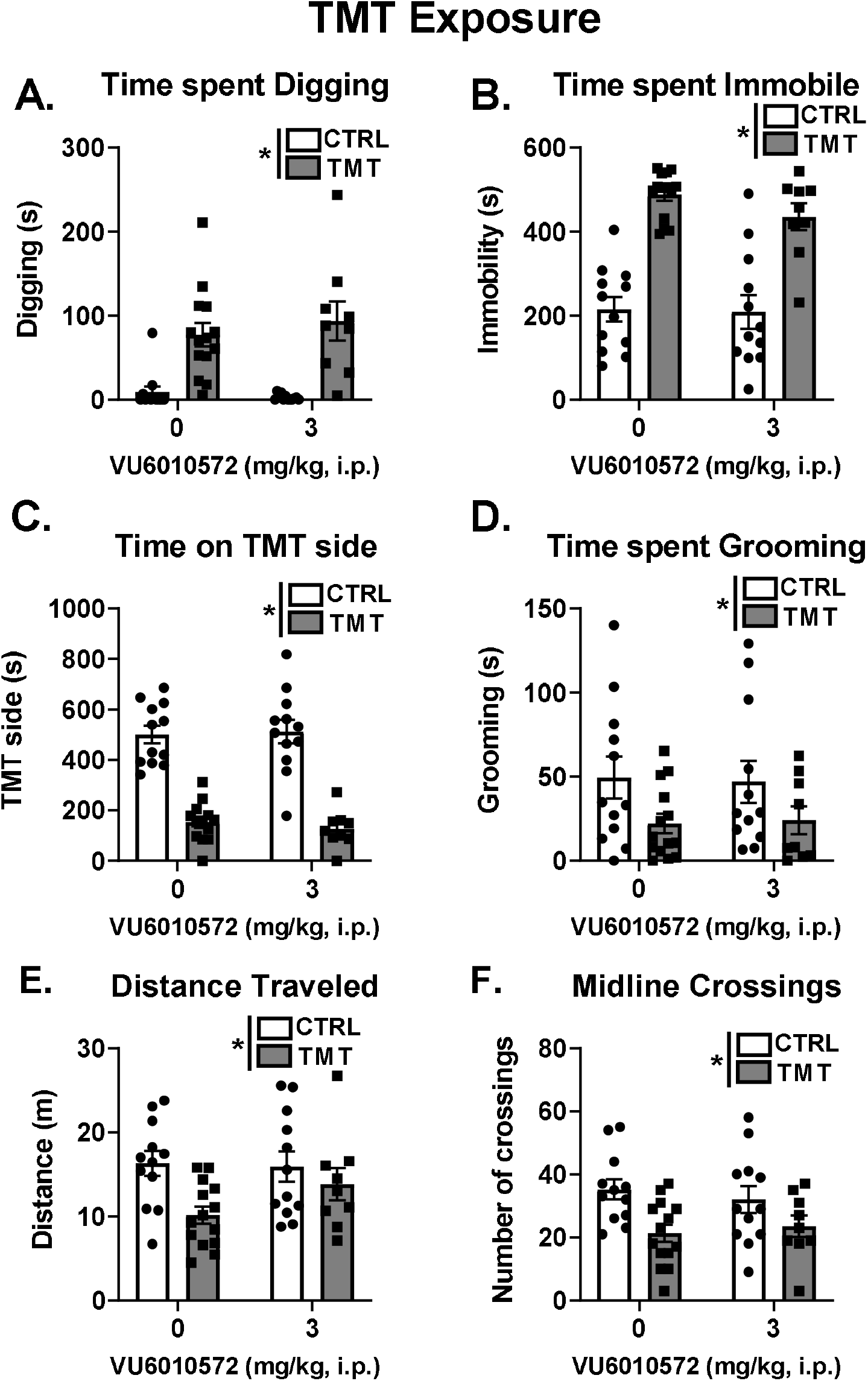
TMT exposure produces stress-reactive behaviors that are not affected by mGlu_3_ NAM. TMT exposure increased time spent digging (A) and immobile (B). TMT exposure decreased time spent on the TMT side (C), time spent grooming (D), distance traveled (E) and the number of midline crossings (F). mGlu_3_ NAM did not affect any of these measures * p≤0.05 significantly different from Control.

### 4.3 TMT exposure and mGlu_3_ NAM produce a hyperactivity phenotype in response to the TMT-paired context

Figure 2 A-D shows the behavioral response in the context in which TMT exposure and mGlu_3_ NAM pretreatment had occurred 2 weeks earlier. For distance traveled (Fig. 2A), a main effect of TMT exposure (F (1, 41) = 13.36, p=0.0007) and mGlu_3_ NAM (F (1, 41) = 9.57, p=0.0036) were observed, both showing increases in distance traveled. No interaction effect was found. For the number of midline crossings (Fig. 2B), a main effect of TMT exposure (F (1, 41) = 11.15, p=0.0018), mGlu_3_ NAM (F (1, 41) = 6.425, p=0.0152) and a TMT x mGlu_3_ NAM interaction effect was observed (F (1, 41) = 5.770. p=0.02). The TMT/vehicle group displayed greater midline crossings compared to the control/vehicle group (p < 0.05). Interestingly, this difference was not observed in the animals that received mGlu_3_ NAM pretreatment. This lack of effect was likely driven by an increase in midline crossings in the control/mGlu_3_ NAM group compared to the control/vehicle group (p < 0.05). For time spent grooming (Fig. 2C), a main effect of TMT showed reduced grooming behavior (F (1, 41) = 5.73, p=0.02), but no significant main effect of mGlu_3_ NAM or interaction was observed. For time spent digging (Fig. 2D), a main effect of TMT exposure was found (F (1, 41) = 4.58, p=0.0382) with increased time digging, but no effect of mGlu_3_ NAM or interaction was observed. No effect of TMT exposure or mGlu_3_ NAM was observed for time spent on the TMT side and time spent immobile (see Table 1). These data indicate that TMT exposure produces hyperactivity in the previously paired TMT context, which may be in part potentiated by mGlu_3_ NAM pretreatment.

**Figure 2.**
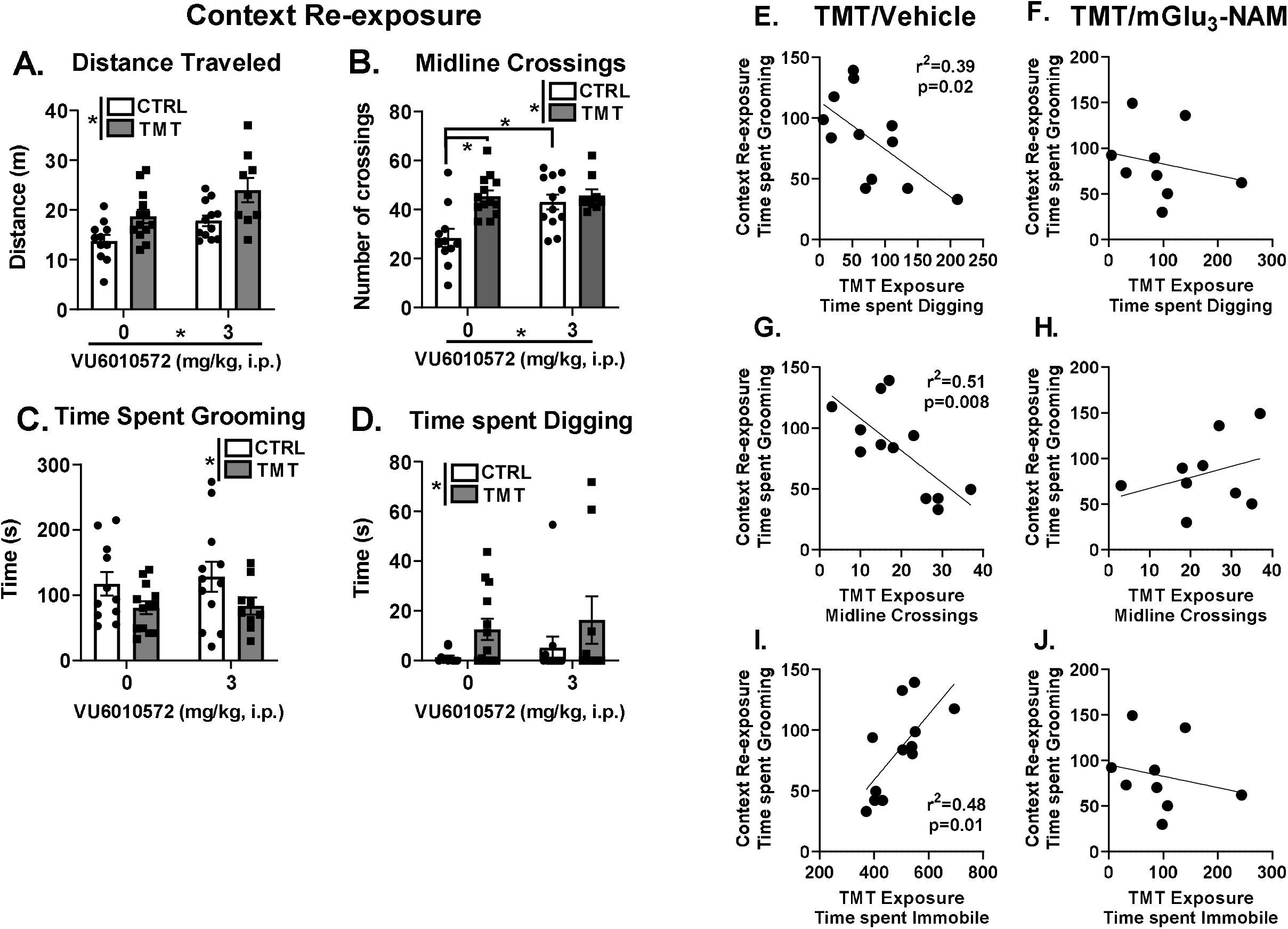
TMT exposure and mGlu_3_ NAM produce a hyperactivity phenotype in response to the TMT-paired context. During the context re-exposure, a main effect of TMT exposure and mGlu_3_ NAM increased both distance traveled (A) and the number of midline crossings (B). Time spent grooming (C) during the context re-exposure was decreased by the TMT exposure. Time spent digging (D) was increased as an effect of TMT exposure. Time spent digging during the TMT exposure negatively correlated with time spent grooming during the context re-exposure in the TMT/vehicle group (E), but not the TMT/mGlu_3_ NAM group (F). The number of midline crossings during the TMT exposure negatively correlated with time spent grooming during the context re-exposure in the TMT/vehicle group (G), but not the TMT/mGlu_3_ NAM group (H). Time spent immobile during the TMT exposure positively correlated with time spent grooming during the context re-exposure in the TMT/vehicle group (I), but not the TMT/mGlu_3_ NAM group (J). * p≤0.05 significantly different from Control

Understanding individual differences in response to stressor may prove valuable in understanding vulnerability to stress-related disorders [61] and behavioral outcomes in rodents [42, 56]. Therefore, correlational analyses were conducted between stress-reactive behaviors during the TMT exposure (Fig. 1) and behavioral responses to the context re-exposure 2 weeks later. Interestingly, in the TMT/vehicle group, time spent grooming during the context re-exposure correlated with 3 distinct stress-reactive behaviors during the TMT exposure. First, time spent digging (Fig. 2E) negatively correlated with time spent grooming (r^2^=0.39, p=0.02). This correlation was not observed in the TMT/mGlu_3_ NAM group (Fig. 2F). The number of midline crossings during the TMT exposure negatively correlated with time spent grooming during the context re-exposure (r^2^=0.51, p=0.008) in the TMT/vehicle group (Fig. 2G), but not in the TMT/mGlu_3_-NAM group (Fig. 2H). Again, time spent immobile during the TMT exposure positively correlated with time spent grooming during the context re-exposure in the TMT/vehicle (Fig. 2I), but not the TMT/mGlu_3_ NAM group (Fig. 2J). These correlations were not observed in either control group (control/vehicle: Digging X Grooming: r=-0.07; Midline Crossings X Grooming: r=-0.11; Immobility X Grooming: r=-0.16; or the Control/mGlu_3_ NAM: Digging X Grooming: r=-0.019; Midline Crossings X Grooming: r=-0.16; Immobility X Grooming: r=-0.13) group. These correlations suggest an association between the degree of stress-reactivity to the TMT exposure with the degree of engagement in grooming behavior during the context re-exposure. Interestingly, these associations were not observed in the TMT-exposed rats that were pretreated with mGlu_3_ NAM suggesting that the mGlu_3_ NAM may disrupt this association.

### 4.4 mGlu_3_ NAM produces lasting anxiolytic-like behavioral effects

Figure 3 shows analysis of the zero-maze test and acoustic startle response test approximately 2 weeks after TMT exposure and mGlu_3_ NAM pretreatment. For percent time spent in the open arms of the zero maze (Fig. 3A), a main effect of mGlu_3_ NAM was found (F (1, 40) = 6.19, p=0.01), indicating greater percent time spent in the open arms, but no effect of TMT exposure or interaction was observed. In the acoustic startle test, there were no significant effects of TMT exposure or mGlu_3_ NAM on average peak startle response (mV, control/vehicle - 770.81 ± 128.30; control/mGlu3 NAM - 957.31 ± 310.94; TMT/Vehicle - 481.33 ± 83.06; TMT/ mGlu_3_ NAM - 1125.47 ± 404.21). However, mGlu_3_ NAM treated rats displayed greater habituation (Fig. 3B) across the 30-trials (F (1, 40) = 4.95, p=0.03), with no significant effect of TMT or interaction. These results indicate that mGlu_3_ NAM produces lasting anxiolytic-like effects and facilitates habituation to the startle response two weeks after the mGlu_3_ NAM injection.

**Figure 3.**
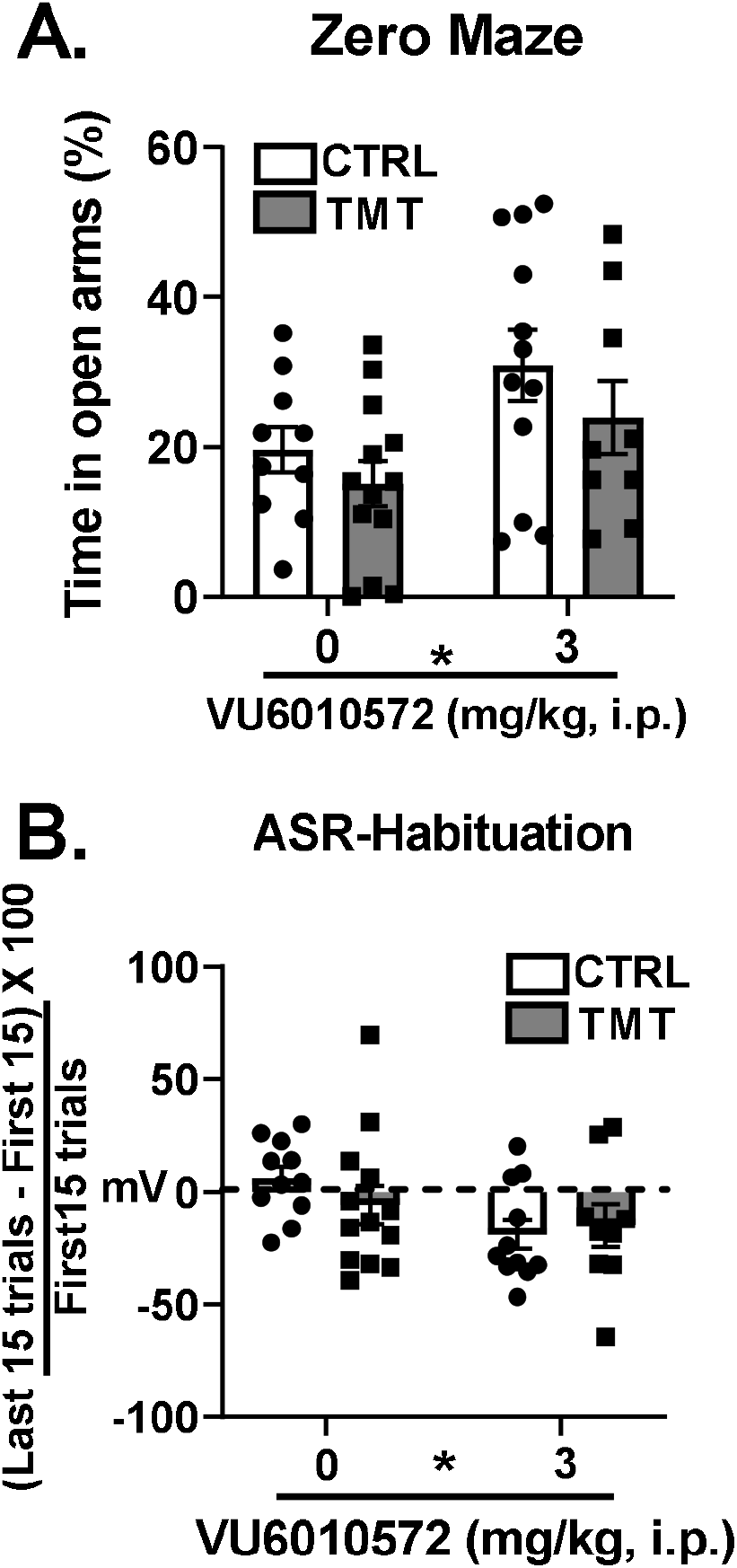
mGlu_3_ NAM produces lasting anxiolytic-like behavioral effects. mGlu_3_ NAM treatment 2 weeks before testing resulted in increased time spent in the open arms of the zero maze test (A) and increased habituation of acoustic startle responses (B). TMT exposure did not affect zero maze or ASR measures. * p≤0.05 significantly different from Control.

### 4.5 TMT exposure produces lasting increases in glutamate receptor gene expression in the insular cortex that are partially mediated by mGlu_3_-signaling during the TMT exposure

Figure 4 shows the effects of TMT exposure and mGlu_3_ NAM on glutamate receptor transcripts in the insular cortex. For AMPA receptor transcripts, *GriA1* (Fig. 4A) and *GriA2* (Fig. 4B) were not affected by TMT exposure or mGlu_3_ NAM. For *GriA3* (Fig. 4C), TMT exposure increased expression (F (1, 23) = 10.54, p = 0.0036), but there was no effect of the mGlu_3_ NAM or significant interaction.

**Figure 4.**
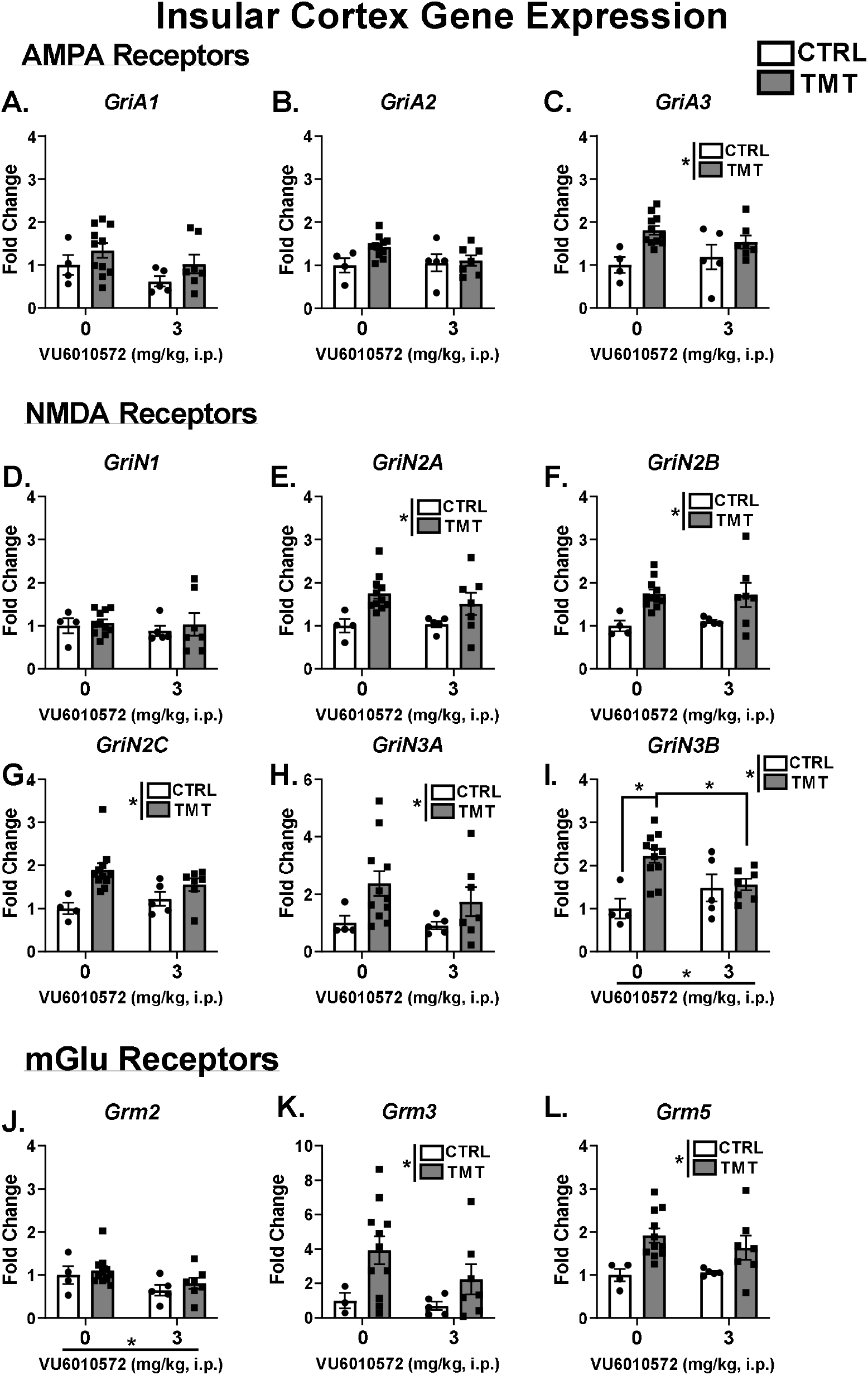
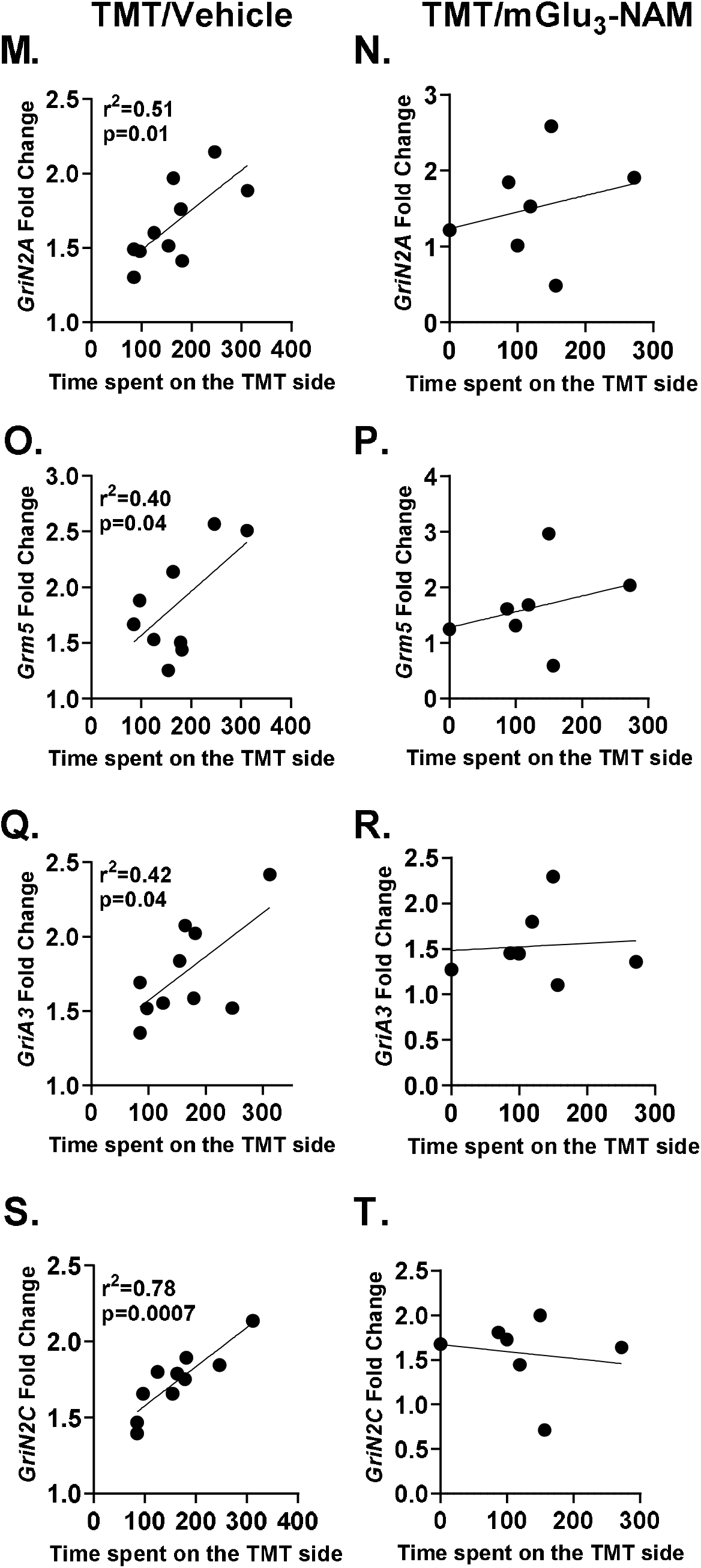
TMT exposure produces lasting increases in glutamate receptor gene expression in the insular cortex that are partially mediated by mGlu_3_-signaling during the TMT exposure. TMT exposure and mGlu_3_ NAM did not affect expression of the AMPA receptor transcripts *GriA1* (A) and *GriA2* (B). The TMT exposure increased expression of *GriA3* (C). TMT exposure and mGlu_3_ NAM did not affect *GriN1* (D) expression, but the TMT exposure increased expression of *GriN2A* (E), *Gri2NB* (F), *GriN2C* (G), *GriN3A* (H) and *GriN3B* (I). *Grm2* (J) was downregulated as an effect of the mGlu_3_ NAM treatment. *Grm3* (K) and *Grm5* (L) were increased by the TMT exposure. In the TMT/vehicle group, but not the TMT/mGlu_3_ NAM group, the amount of time the animal spent on the side of the chamber containing the TMT odor was positively correlated with *GriN2A* (M, N), Grm5 (O, P), *GriA3* (Q, R) and *GriN2C* (S, T), and expression in the insular cortex. * p≤0.05 significantly different from Control.

For NMDA receptor transcripts, no effect of TMT exposure or mGlu_3_ NAM was found for *GriN1* expression (Fig. 4D). For *GriN2A* (Fig. 4E), *GriN2B* (Fig. 4F), *GriN2C* (Fig. 4G), and *GriN3A* (Fig. 4H), TMT exposure increased expression (*GriN2A* - F(1, 23) = 10.24, p = 0.004), *GriN2B* - F (1, 23) = 13.15, p = 0.0014), *GriN2C* - F (1, 23) = 11.53, p = 0.0025, *GriN3A* - F (1, 23) = 5.001, p = 0.03) but did not show an effect of the mGlu_3_ NAM or significant interaction. No effect of TMT exposure, mGlu_3_ NAM or significant interaction was observed for *GriN2D* (fold changes; control/vehicle - 1.0 ± 0.15; control/ mGlu_3_ NAM - 1.27 ± 0.33; TMT/Vehicle - 1.82 ± 0.17; TMT/mGlu3 NAM - 1.26 ± 0.16). *GriN3B* expression (Fig. 4I) was increased by TMT exposure (F (1, 23) = 9.13, p = 0.006), but not affected by mGlu_3_ NAM, and a significant TMT exposure x mGlu_3_ NAM interaction was observed (F (1, 23) = 7.17, p= 0.01). Post-hoc analyses showed that *GriN3B* expression was increased in the TMT/vehicle group compared to the control/vehicle group (p < 0.05), an effect that was prevented by mGlu_3_ NAM pretreatment (i.e., no difference between TMT/ mGlu_3_ NAM vs. control/mGlu_3_ NAM groups). Additionally, *GriN3B* was decreased in the TMT/mGlu_3_ NAM group compared to the TMT/vehicle group (p < 0.05), but not in the control/mGlu_3_ NAM compared to control/vehicle groups (p > 0.05). These data indicate that mGlu_3_-signaling during TMT exposure mediates the stressor-induced increase of *GriN3B* in the insular cortex.

For mGlu receptor transcripts, *Grm2* (Fig. 4J) was not affected by the TMT exposure, but was downregulated as a main effect of mGlu_3_ NAM (F (1, 23) = 5.19, p = 0.03). *Grm3* (Fig. 4K) and *Grm5* (Fig. 4L) were upregulated as a main effect of TMT exposure (*Grm3* - F (1, 22) = 5.384, p = 0.03; *Grm5* – F (1, 23) = 11.48, p=0.0025), with no effect of the mGlu_3_ NAM or significant interaction. Together, these data suggest that excitatory adaptations in the insular cortex as a lasting result of TMT exposure is partially mediated by mGlu_3_-signaling during the stressor.

Examination of associations between behavioral reactivity during the TMT exposure and gene expression showed that in the vehicle-treated rats in the TMT group, time spent on the TMT side of the TMT exposure chamber positively correlated with several of the upregulated glutamate receptors in the insular cortex. Time spent on the TMT side positively correlated with *GriN2A* in the TMT/vehicle group (Fig. 4M; r=0.71, p=0.02), but not in the TMT/mGlu3-NAM group (Fig. 4N). Time spent on the TMT side positively correlated with *Grm5* (Fig. 4O; r=0.63, p=0.04) in the TMT/vehicle group, but not the TMT/mGlu_3_ NAM group (Fig. 4P). Time spent on the TMT side positively correlated with *GriA3* (Fig. 4Q; r=0.65, p=0.04) in the TMT/vehicle, but not the TMT/mGlu_3_ NAM group (Fig. 4R). Finally, time spent on the TMT side positively correlated with *GriN2C* (Fig. 4S; r=0.88. p=0.001) in the TMT/vehicle group, but not in the TMT/mGlu_3_ NAM group (Fig. 4T). These results indicate that proximity to the TMT exposure during the stressor is associated with the lasting glutamate receptor upregulation in the insular cortex. Interestingly, none of these correlations were observed in the TMT/mGlu_3_ NAM group, indicating that mGlu_3_ NAM pretreatment disrupted this association. Neither control groups showed these correlations either: control/vehicle (Time on TMT side during exposure by: *GriN2A* – r=-0.73; *GriN2C* – r=-0.61; *GriA3* – r=-0.47; *Grm5* – r=-0.34) or control/ mGlu_3_ NAM (Time on TMT side during exposure by: *GriN2A* – r=-0.15; *GriN2C* – r=-0.41; *GriA3* – r=-0.50, *Grm5* – r=0.40) group. No other significant correlations were observed in any group.

### 4.6 mGlu_3_ NAM upregulates NMDA receptor subunits in the BNST, but this effect is prevented by TMT exposure

Figure 5 shows the gene expression changes in the BNST following TMT exposure and mGlu_3_ NAM pretreatment. For *GriN1* (Fig. 5A), no effect of TMT or mGlu_3_ NAM was found. For *GriN2A* (Fig. 5B), there was a main effect of TMT exposure (F (1, 21) = 8.24, p = 0.0091), no main effect of mGlu_3_ NAM, but a significant interaction effect (F (1, 21) = 17.18, p = 0.0005). mGlu_3_ NAM increased *GriN2A* expression in controls (p < 0.05) but not in the TMT group. *GriN2A* expression was decreased by mGlu_3_ NAM in the TMT group (p < 0.05), but not changed in vehicle-treated rats. For *GriN2B* (Fig. 5C), there was a significant main effect of TMT exposure (F (1, 21) = 5.43, p = 0.02), a main effect of mGlu_3_ NAM (F (1, 21) = 6.25, p = 0.02), and a significant interaction effect (F (1, 21) = 7.44, p = 0.01). mGlu_3_ NAM increased expression in controls (p < 0.05) but not in the TMT group. Again, TMT exposure decreased expression in mGlu_3_ NAM treated (p < 0.05), but not in vehicle treated rats (p > 0.05). Post-hoc tests did not show any significant effects of mGlu_3_ NAM in either group. For *GriN3B* (Fig. 5D), no main effect of TMT or mGlu_3_ NAM was observed, but a significant interaction effect was found (F (1, 21) = 11.97, p = 0.0023). Again, mGlu_3_ NAM increased *Grin3B* expression in controls (p < 0.05) but not in the TMT group (p > 0.05). TMT exposure decreased expression in mGlu_3_ NAM treated (p < 0.05), but not in vehicle treated rats. For *Grm2* (Fig. 5E), a main effect TMT (F (1, 21) = 6.16, p = 0.02) was observed, no effect of mGlu_3_ NAM, but a significant TMT x mGlu_3_NAM interaction effect (F (1, 21) = 5.31, p = 0.03) was observed. Post hoc effects of TMT exposure showed decreased expression in rats treated with mGlu_3_ NAM (p < 0.05), but not in vehicle-treated rats (p > 0.05). *Grm5* (Fig. 5F) expression was not affected by TMT exposure or mGlu_3_ NAM. For *GriN3A, GriN2C, GriN2D, GriA1, GriA2, GriA3*, and *Grm3* no effect of TMT, mGlu_3_ NAM or interaction were observed (Table 3). These data indicate that mGlu_3_ NAM increased expression of NMDA receptor sub-units in controls, but TMT exposure prevented this upregulation, suggesting differential transcriptional regulation by mGlu_3_-signaling in the BNST that is dependent on stressor conditions.

**Figure 5–.**
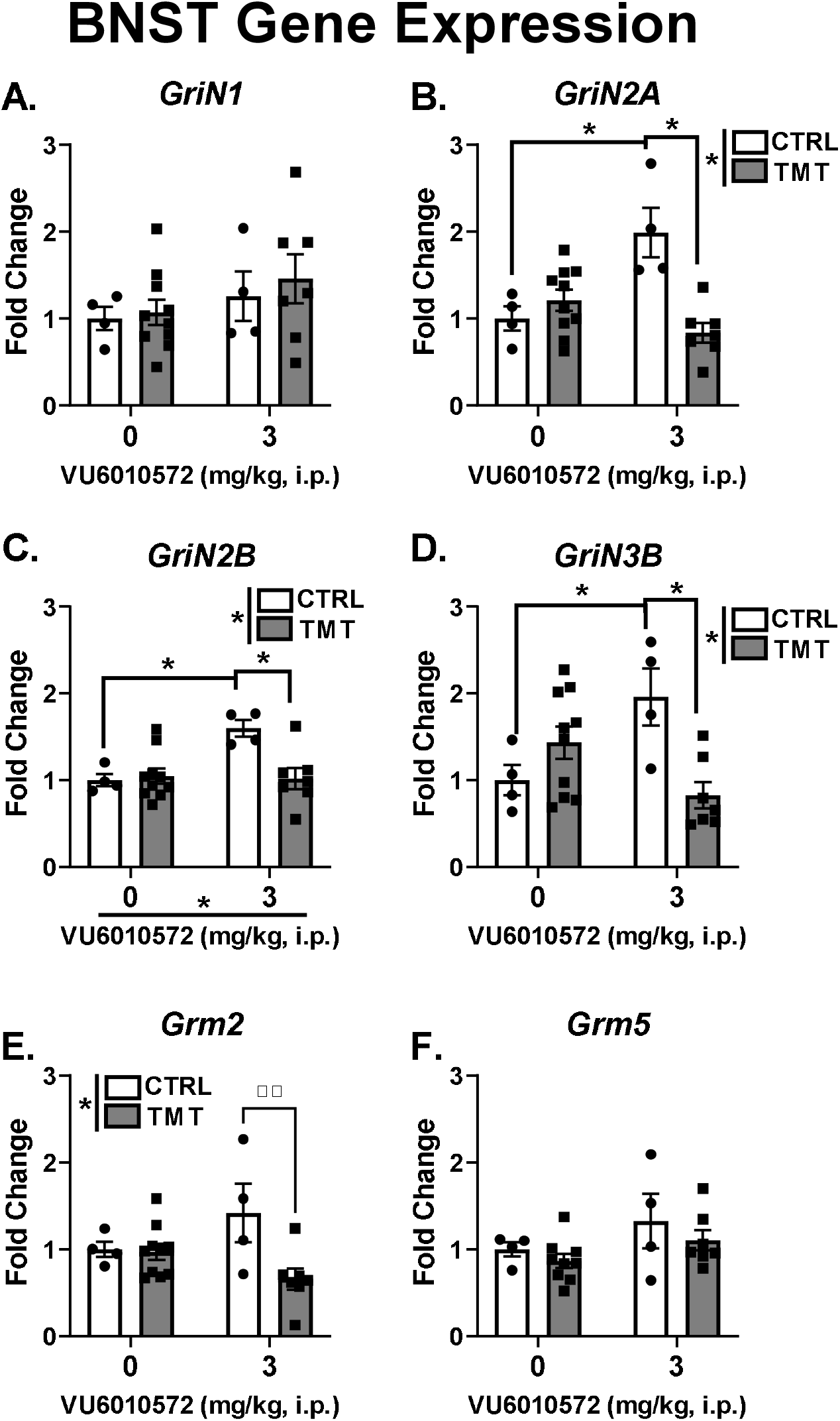
mGlu_3_ NAM upregulates NMDA receptor subunits in the BNST, but this effect is prevented by TMT exposure. The TMT exposure and mGlu_3_ NAM did not affect *GriN1* (A) gene expression in the BNST. *GriN2A* (B), *GriN2B* (C) and *GriN3B* (D) were increased by mGlu_3_ NAM, but these effects were not observed in rats exposed to TMT. For *Grm2* (E), TMT exposure resulted in decreased expression in the mGlu3 NAM treated group only. *Grm5* (F) was not affect by TMT exposure or mGlu3 NAM.

**Table 2 –.** Context Re-exposure –. Table 2 shows the context re-exposure behavioral measures that were not reported in the results section in seconds of the total 15 min test.

| Context Re-exposure Figure 3 – Mean ± SEM | | |
| --- | --- | --- |
| Group | Time spent Immobile (s) | Time spent on TMT side (s) |
| CTRL/Veh | 163.3 ± 18.8 | 131.4 ± 14.8 |
| CTRL/mGlu<sub>3</sub> | 137.0 ± 10.2 | 169.8 ± 9.8 |
| TMT/Veh | 141.9 ± 17.8 | 158.9 ± 7.8 |
| TMT/mGlu<sub>3</sub> | 117.2 ± 11.3 | 165.0 ± 7.7 |

**Table 3 –.**
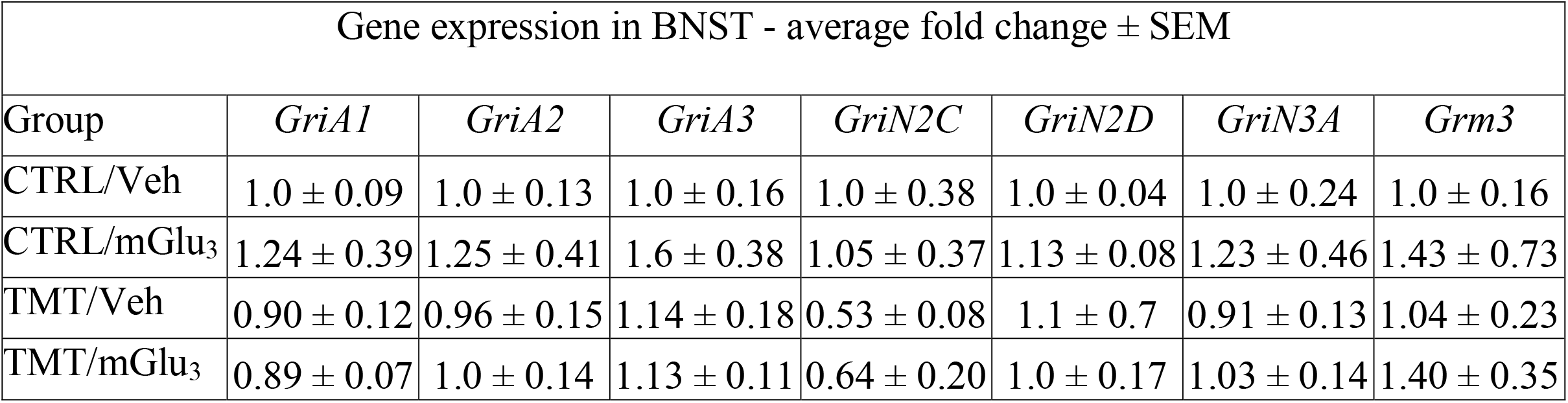
Gene expression in the BNST –. Table 3 shows the gene expression (Fold change of control/vehicle group) results in the BNST.

| Gene expression in BNST - average fold change ± SEM | | | | | | | |
| --- | --- | --- | --- | --- | --- | --- | --- |
| Group | *GriA1* | *GriA2* | *GriA3* | *GriN2C* | *GriN2D* | *GriN3A* | *Grm3* |
| CTRL/Veh | 1.0 ± 0.09 | 1.0 ± 0.13 | 1.0 ± 0.16 | 1.0 ± 0.38 | 1.0 ± 0.04 | 1.0 ± 0.24 | 1.0 ± 0.16 |
| CTRL/mGlu<sub>3</sub> | 1.24 ± 0.39 | 1.25 ± 0.41 | 1.6 ± 0.38 | 1.05 ± 0.37 | 1.13 ± 0.08 | 1.23 ± 0.46 | 1.43 ± 0.73 |
| TMT/Veh | 0.90 ± 0.12 | 0.96 ± 0.15 | 1.14 ± 0.18 | 0.53 ± 0.08 | 1.1 ± 0.7 | 0.91 ± 0.13 | 1.04 ± 0.23 |
| TMT/mGlu<sub>3</sub> | 0.89 ± 0.07 | 1.0 ± 0.14 | 1.13 ± 0.11 | 0.64 ± 0.20 | 1.0 ± 0.17 | 1.03 ± 0.14 | 1.40 ± 0.35 |

## 5. Discussion

These studies show that mGlu_3_-signaling during exposure to the TMT predator odor stressor contributes to both behavioral (context reactivity) and molecular (glutamate receptor gene expression) adaptations approximately 2 weeks after the stressor exposure in male rats. Rats engaged in stress-reactive behaviors during the TMT exposure, and a hyperactive behavioral phenotype during the context re-exposure (in the absence of TMT) two weeks later, which may reflect hypervigilance to the stressor context [58]. TMT exposure did not produce differences in the zero maze or ASR test, but a single administration of the mGlu_3_ NAM VU6010572 produced lasting anxiolytic-like effects on the zero maze test and promoted greater habituation in arousal responses in the ASR test. Examination of gene expression in the insular cortex 17 days after the TMT exposure, showed upregulation of mGlu, NMDA and AMPA receptor transcripts as a main effect of TMT exposure, indicating excitatory adaptations to stressor. Blocking mGlu_3_ receptor signaling with VU6010572 (mGlu_3_ NAM pretreatment) before TMT exposure restored *GriN3B* expression to control/vehicle levels in the insular cortex, reflecting a role for mGlu3 signaling during TMT exposure on the lasting NMDA receptor adaptations in the insular cortex. In contrast to the insular cortex, glutamate receptor gene expression in the BNST was more sensitive to mGlu_3_ NAM than to the TMT exposure. mGlu_3_ NAM pretreatment increased NMDA receptor expression (*GriN2A, GriN2B, GriN3B*), which was prevented/restored by the TMT exposure. This indicates NMDA receptor transcriptional regulation in the BNST by mGlu_3_-signaling that is sensitive to stress conditions.

Consistent with previous findings, the TMT exposure produced engagement in stress-reactive behaviors (immobility, digging, avoidance, and decreased grooming, distance traveled and midline crossings), indicating that TMT exposure is an acute stressor [56, 58]. Pretreatment with the mGlu_3_ NAM (VU6010572) did not affect any of these behaviors, suggesting that mGlu_3_-signaling during the stressor does not contribute to the engagement of acute stress-reactive behaviors. Prior studies in mice show that a similar mGlu_3_ NAM (VU0650786) administered prior to restraint stress prevented decreased sucrose intake indicative of motivational behavior caused by the restraint stress [19] and that VU6010572 produced acute anti-depressant-like behavioral effects [29]. Together with the present findings, these data serve to specify the acute effects of mGlu_3_ NAM, which may be related more to anti-depressant-like effects rather than effects on stress-reactivity.

Similar to previous reports from our lab, rats exposed to TMT showed hyperactivity (increased distance traveled, midline crossings and digging, and decreased grooming) when returned to the TMT context in the absence of TMT 2 weeks later [58]. These behavioral changes may reflect a hypervigilance phenotype in response to the stressor context as previously proposed [58]. Correlational analyses in the TMT/vehicle group showed that time spent grooming during the context re-exposure negatively correlated with both time spent digging and the number of midline crossings during the TMT exposure. Additionally, time spent immobile during the TMT exposure positively correlated with time spent grooming during the context re-exposure. These data patterns suggest a potential association between stress-reactivity (digging, mid-line crossings, immobility) during the TMT exposure with the lasting behavioral adaptations in response the context re-exposure (i.e. diminished grooming). Interestingly, these correlations in the TMT/vehicle group were not observed in the TMT/mGlu_3_ NAM group, suggesting that mGlu_3_ NAM pretreatment prior to stressor disrupts the associations between TMT exposure and context re-exposure stress reactivity measures. Surprisingly, while mGlu_3_ NAM pretreatment had no effect on stress-reactivity during the TMT exposure, mGlu3 NAM pretreatment increased distance traveled and midline crossings during the context re-exposure similar to the effect of TMT exposure. Furthermore, the increase in midline crossings was only observed in the vehicle-treated rats, but not in rats pretreated with mGlu_3_ NAM, driven by an increase in midline crossings in the control/mGlu_3_ NAM group compared to the control/vehicle group. This data pattern suggests that the mGlu_3_ NAM may contribute to the hypervigilance phenotype produced by TMT stressor, and/or a lasting increased locomotion/activity phenotype in general. Prior studies show that TMT exposure decreases gene expression of *Grm3* (encodes for mGlu_3_ receptors) [58]; therefore, these results suggest the possibility that mGlu_3_ loss of function, either through pharmacological targeting (mGlu_3_ NAM) or stressor [19] (TMT exposure), could result in increased locomotion/activity associated with the stressor-induced context reactivity phenotype.

The mGlu_3_ NAM pretreatment also produced other lasting behavioral changes. While TMT exposure did not affect zero maze and ASR measures, the mGlu_3_ NAM produced anxiolytic effects on the zero maze and promoted greater habituation of arousal response in the ASR test. Because mGlu_3_ NAM was administered 2 weeks prior to behavioral tests, these findings suggest a lasting effect of the mGlu_3_ NAM on anxiety and arousal processes, which may contribute to the therapeutic potential of negative allosteric modulation of mGlu_3_ as to date VU6010572 has been shown to have rapid anti-depressant-like effects [29]. Further, mGlu_2/3_ antagonists have been proposed as potential alternative anti-depressants with similar acute and lasting efficacy and mechanisms to ketamine in the absence of abuse liability [17]. Therefore, these novel findings on anxiety-like behavior and arousal habituation show that a single administration of the mGlu_3_ NAM VU6010572 is sufficient to produce lasting therapeutic effects, and therefore may have potential as a new glutamate targeting therapeutic.

TMT exposure produced lasting increases of several glutamate receptor transcripts in the insular cortex, including mGlu (*Grm5*), NMDA (*GriN2A, GriN2B, GriN2C, GriN3A, GriN3B*) and AMPA (*GriA3*) receptors. Because many of these genes encode for synaptic glutamate receptors [62], these data may reflect changes to excitatory synaptic number and/or density. Alternatively, these data could reflect differences in the composition of glutamate receptors that potentially result in functional differences. Because the insular cortex is involved in negative affect, anxiety, fear memories and threat conditioning [31, 34, 35, 41, 63], and NMDA receptor adaptations are involved in synaptic plasticity in the insular cortex [64], these adaptations could in part underlie the hypervigilance phenotype observed in the context re-exposure test.

Interestingly, mGlu_3_ NAM treatment prior to TMT exposure completely reversed the upregulation of *GriN3B* (encodes GluN3B) induced by TMT exposure. GluN3B is an NMDA receptor subunit expressed at low levels in the brain [65] but may play an important role in the lasting alterations from synaptic plasticity and synaptogenesis [66]. In contrast to canonical NMDA receptors (GluN1-GluN2 heterotetramers), GluN3-containing NMDA receptors (GluN1-GluN3A/B) have a smaller single channel conductance, lower Ca2+ permeability, less sensitivity to the Mg2+ block, and are located outside of the synapse [66]. Early studies indicated that the GluN3 subunits diminished glutamate-evoked NMDA currents [67], which was proposed to indicate a dominant-negative function of GluN3B on traditional GluN1-GluN2 currents. Therefore, these data could indicate that mGlu_3_-signaling during the TMT predator odor stressor diminishes synaptic/phasic NMDA receptor function through sequestration of glutamate by the dominant-negative sub-unit GluN3B. Interestingly, GluN1-GluN3 heterotetramers are not sensitive to glutamate or NMDA, but all 4 subunits bind glycine [66]. Because endogenous concentrations of glycine constitutively occupy these receptors, it has been proposed that GluN1/GluN3 heterotetramers remain open *in vivo* and serve to modulate tonic excitation [66, 68]. Therefore, the upregulation of *GriN3B* transcript expression in the insular cortex by stressor-induced mGlu_3_-signaling might also reflect enhanced tonic excitation in the insular cortex as a lasting adaptation to stressor. As such, important future directions will examine how these adaptations affect cellular physiology in the insular cortex. Importantly, these findings are the first to show that TMT exposure produces lasting adaptations to excitatory gene expression in the insular cortex, and that mGlu_3_-signaling during stressor mediates *GriN3B* transcriptional upregulation.

Finally, the mGlu_3_-NAM decreased *Grm2* (encodes for mGlu_2_) expression in the insular cortex, indicating that mGlu_3_ signaling is involved in the transcriptional regulation of *Grm2* in the insular cortex. This adaptation might, in part, underlie the anxiolytic and arousal habituation effects of mGlu_3_ NAM as mGlu2 is involved in maladaptive behavioral responses to stress [27].

Correlational analyses showed that time spent on the TMT side of the test chamber during the TMT exposure positively correlated with the expression of *GriN2A, GriN2C, GriA3*, and *Grm5* in the insular cortex in the TMT/Vehicle group. Therefore, rats that spent more time near the TMT stressor showed higher upregulation of glutamate receptors in the insular cortex. Similar to the correlations observed between stress-reactive behaviors to TMT exposure and the context re-exposure reactivity measures, correlations between time spent on the TMT side and insular cortex glutamate receptor gene expression were not observed in the TMT/mGlu_3_ NAM (or control) group. These findings suggest the possibility that mGlu_3_ NAM pretreatment disrupted the relationship between time spent near the TMT source and increased expression of *GriN2A, GriN2C, GriA3*, and *Grm5* in the insular cortex.

While the insular cortex was sensitive to the lasting effects of TMT exposure, glutamate receptor gene expression in the BNST was more response to the mGlu_3_ NAM. Treatment with the mGlu_3_ NAM resulted in the upregulation of some NMDA receptor subunit transcripts (*GriN2A, GriN2B* and *GriN3B*) in the control/mGlu_3_ NAM group compared to the control/vehicle group. Interestingly, these of mGlu_3_ NAM effects were not observed in rats exposed to TMT. For *Grm2*, TMT exposure downregulated expression in the mGlu_3_-treated group but not in the vehicle group. First, these data show that two weeks following a single mGlu_3_ NAM pretreatment there are changes in excitatory gene expression in the BNST, which may in part underlie its therapeutic effects on zero maze and ASR habituation. Failure to observe similar increases in the TMT/mGlu_3_ NAM group indicates that this transcriptional regulation is sensitive to stress conditions, and/or that the mGlu_3_ NAM produces different effects on transcription of glutamate receptors based on stress conditions. Furthermore, prior data suggests that mGlu_3_ receptors may be activated during the TMT exposure [19, 58]. Therefore, the upregulation of these NMDA receptor transcripts induced by blocking mGlu_3_ may be offset by the subsequent activation of mGlu_3_ induced by the TMT exposure, thus preventing the upregulation of the NMDA receptor transcripts. Indeed, prior studies have also demonstrated involvement of mGlu_2/3_ signaling in stress-induced adaptations to the BNST. Specifically, mGlu_2/3_ agonism (LY379268) potentiated foot-shock induced c-Fos increases in the dorsolateral BNST [69]. Therefore, because the BNST is known for its involvement in lasting adaptation to stress [38, 70], these NMDA receptor adaptations could in part underlie the lasting anxiolytic and ASR habituation effects of mGlu_3_ NAM.

The insular cortexr cortex and BNST form reciprocal projections [71, 72] and both regions contribute to affective behavioral measures in rodents [34, 35, 73] and humans [31]. Recent studies have shown that insular cortex/BNST circuitry are involved in negative affective measures associated with alcohol withdrawal in rodents [41]. Therefore, these findings showing stressor- and mGlu_3_ NAM -induced adaptations to glutamate receptors in the insular cortex and BNST could reflect adaptations to insular cortex/BNST excitatory circuitry as a result of stressor and mGlu_3_ NAM, and further implicate this circuitry in stress-induced behavioral changes. Future work will investigate the functional involvement of insular cortexr cortex/BNST circuitry on lasting behavioral adaptations of the TMT exposure (i.e. context re-exposure hyperactivity and anxiolytic effects of mGlu_3_ NAM).

An important consideration for the present findings is that like the TMT groups, the control groups experienced exposure to a novel environment (though in the absence of TMT). This experience is potentially a stressor and therefore, this feature of our experimental design should be considered when making conclusions about the effects of mGlu_3_ NAM and the observed adaptations in behavior and gene expression. Furthermore, the effects of TMT exposure and mGlu_3_-signaling were not evaluated in female rats, which is a limitation to these studies as sex differences may exist in response to TMT stressor and mGlu_3_ NAM [56]. Furthermore, the BNST is sexually dimorphic [74], suggesting the possibility of sex differences in the gene expression changes in the BNST specifically. Important follow-up studies will evaluate these effects in female rats.

These findings show that mGlu_3_ NAM did not affect acute stress-reactivity in response to TMT exposure, but did contribute to lasting phenotypes induced by the TMT exposure. TMT exposure produced a hypervigilant/hyperactive phenotype in response to the context re-exposure 2 weeks later, and mGlu_3_ NAM pretreatment potentiated some of the hyperactive behavioral measures. Gene expression analysis show that TMT exposure produced adaptations to glutamate receptors in the insular cortex, and that mGlu_3_ NAM during stressor mediated the transcriptional upregulation of *GriN3B*. Unlike the data pattern in the insular cortex, gene expression in the BNST was not affected by TMT exposure under vehicle conditions, but rather TMT exposure prevented the robust effects of mGlu_3_ NAM on NMDA receptor upregulation in the BNST. Finally, mGlu_3_ NAM produced lasting anxiolytic-like behavioral phenotypes. These findings show that mGlu_3_ receptors are involved in lasting adaptations to stressor may reflect a promising target for stressor-related neuropsychiatric disorders.

## Acknowledgements

This work was supported in part by the National Institute of Health AA026537 and AA011605 (JB) and by the Bowles Center for Alcohol Studies. RET and MB were supported in part by NS007431.

## Conflicts of interest

none.

## Data Availability Statement

Data available on request from the authors.

## Notes

### Competing Interest Statement

The authors have declared no competing interest.

